# Vasculature-staining with lipophilic dyes in tissue-cleared brains assessed by deep learning

**DOI:** 10.1101/2020.05.16.099705

**Authors:** Beatriche L. E. Henriksen, Kristian H. R. Jensen, Rune W. Berg

## Abstract

Visualization of the vasculature in three dimensions (3D) has become attractive, particularly in stroke models. 3D-reconstruction is aided by tissue-clearing, where the transparency allows imaging of fluorescent probes in deeper structures. The vasculature is commonly stained by fluorescent lipophilic dyes that are incorporated into the wall during transcardial perfusion. Nevertheless, tissue clearing involves extracting the light-scattering lipids, and hence also the lipid-appended dyes. The wash-out likely depends on dye and its aldehyde-fixability. Fixation secures cross-linking to proteins and hence retainment in the tissue. However, the compatibility of various types of dyes is largely unknown. We tested and compared 9 different dyes for vasculature staining and tolerance to lipid clearing, which was quantified using deep learning image segmentation. Among the dyes, we found a subset that is both cost-effective and compatible with tissue lipid clearing. We suggest these dyes will provide a valuable tool for future investigations.

**Highlights:**

- Size, alkyl-chains and aldehyde-fixable groups improve dye retention and performance.
- Cost-effective dyes in specific liposomes result in optimal vessel staining.
- SP-DiIC18 is compatible with CLARITY and BrainFilm and advantageous for most studies.
- We recommend, SP-DiI and R18, cost-effective dyes, for vessel-painting with CLARITY.

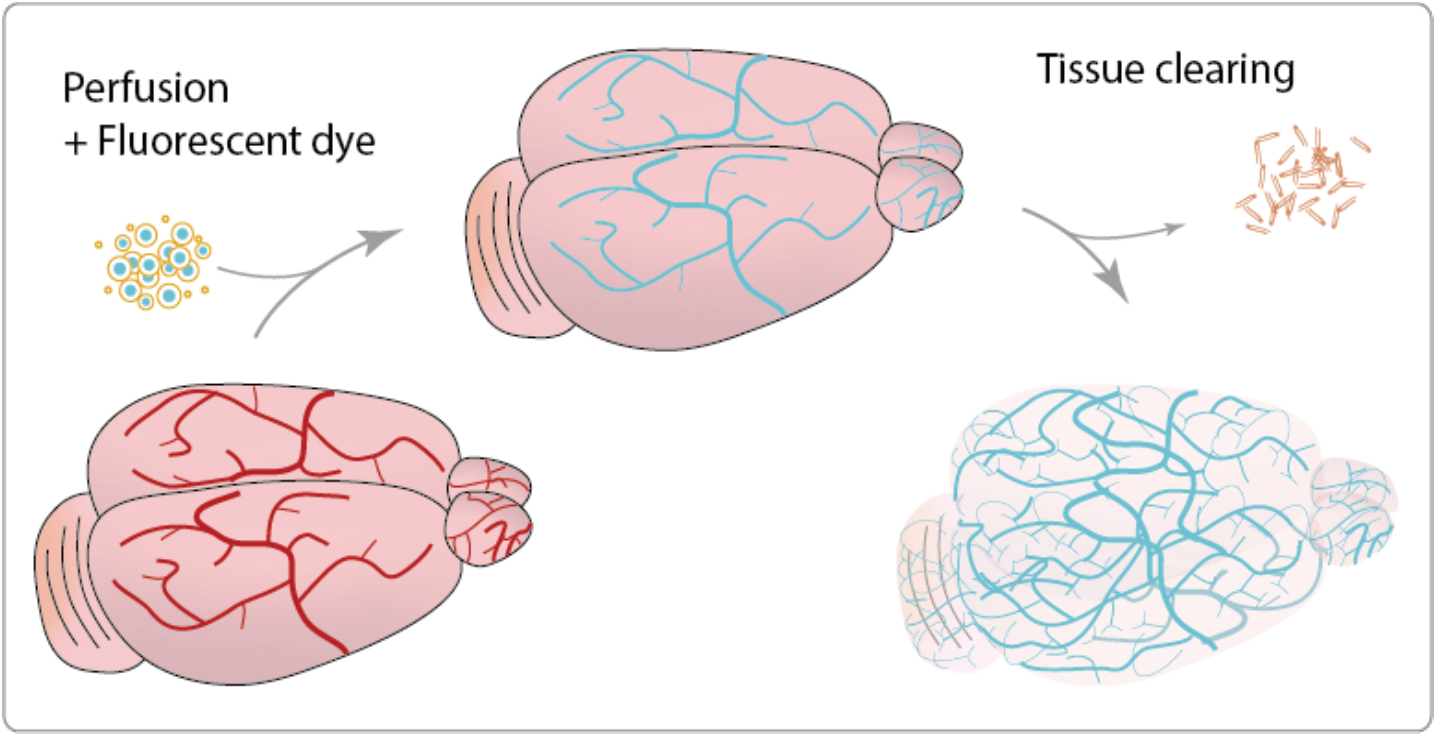

## Introduction

Blood flow is important in basic research as well as in pathology such as stroke, and the reconstruction of the vasculature in three-dimensions (3D) is a valuable investigative tool. Within the last decade several tissue-clearing techniques, have been developed, which make large samples transparent and enables microscopy of the 3D structures within intact organs (Ertürk et al., 2012; Chung et al., 2013; Yang et al., 2014; Epp et al., 2015; Zheng and Rinaman, 2016; Kolesová et al., 2016; Neckel et al., 2016; Liu et al., 2017; Carrillo et al., 2018; Qi et al., 2019; Cai et al., 2019). One of the early, and still, powerful methods is CLARITY (Clear Lipid-exchanged Acrylamide-hybridized Rigid Imaging-compatible Tissue-hYdrogel)(Gradinaru et al., 2018; Jensen and Berg, 2017), which utilizes an acrylamide-hydrogel to keep proteins bonded in a skeleton structure and the removal of lipids, which are the primary cause of light-scattering in tissues, by using a detergent leaving the tissue transparent (Chung et al., 2013; Tomer et al., 2014).

Another challenge is to visualize specific entities of interest, such as neurons, glia, or vasculature. The endogenous expression of fluorescent proteins is a solution in cells, however, since this can be difficult in some cases, it is beneficial to be able to deliver a fluorescent dye to the specified target, which can be retained after tissue-clearing. With exogenous fluorescent molecules, it is possible to stain specific targets, such as tracking the location of electrodes or axonal tracing (Jensen and Berg, 2016). The vasculature can be stained using similar dyes by perfusion, and the tissue can then be cleared using CLARITY, leaving the vasculature visible in 3D (Li et al., 2008; Konno et al., 2017; Salehi et al., 2018).

Nevertheless, lipophilic dyes are likely to be removed unintentionally when clearing the tissue of lipids. Modified lipophilic dyes, with membrane binding or aldehyde fixable side chains, may work better. So far, only few lipophilic dyes, as well as vessel staining methods, have been tested in large, CLARITY-cleared tissue samples (Chung et al., 2013; Jensen and Berg, 2016). Several CLARITY-compatible lipophilic dialkylcarbocyanine (DiI) dye analogs have been identified for electrode and axonal tracing (Jensen and Berg, 2016). DiI dyes are known for inserting their aliphatic chains into the lipid bilayer of cell membranes (Axelrod, 1979), thus anchoring the dye to lipids.

We tested the compatibility of nine lipophilic dyes with perfusion vessel staining and lipid clearing (figure 1): DiIC18 and the sorter DilC12 that are well characterized and previously used in vessel staining in thin sections (Konno et al., 2017), in addition to three fixable variants of these, the CM-DiIC18, SP-DiIC18, and DiIC12-DS with increased solubility and resistance to detergents. We also tested four other lipophilic dyes, FM 1-43FX, Nile Red, Octadecyl Rhodamine B (R18) and BODIPY TR Methyl Ester. Dye variants with red emission were selected for the most straightforward comparison.

**Figure 1:**
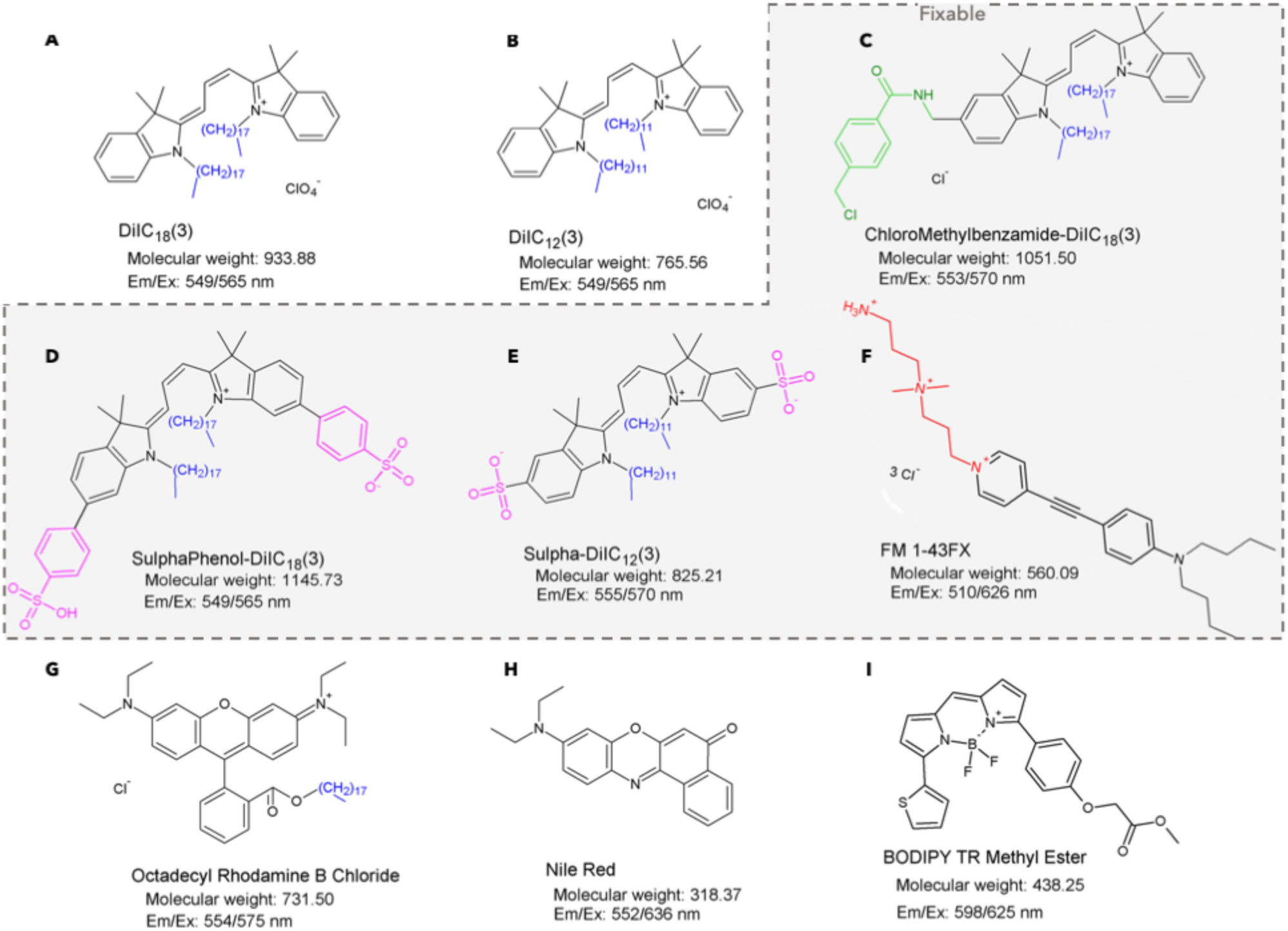
Lipophilic dyes tested for CLARITY-compatibility. The cationic indocarbocyanine dye, DiIC18 (A) and the shorter 12-carbon analog DiIC12 (B) both rely on their alkyl-tails (blue) to retain in the lipid bilayer. However, DiIC12 diffuses more rapidly with better membrane incorporation (Mukherjee et al., 1999) and slower aggregation time in solution (Konno et al., 2017). We selected three aldehyde-fixable analogs: CM-DiIC18 (C) with a thiol-reactive chloromethylbenzamido (CM) group (green), SP-DiIC18 (D) with two sulfophenyl (SP) and DiIC12-DS (E) with two directly sulfonated (DS) groups (magenta). G) FM 1-43FX is a styryl dye with an aliphatic amine side chain for aldehyde fixability (red), which binds rapidly to the plasma membrane and has a strong fluorescent signal when bound (Bolte et al., 2004). G) Octadecyl Rhodamine B (R18) is Rhodamine B attached to a single 18-carbon alkyl-tail. H) Nile Red (commonly used in histology to stain cells) is a generic lipophilic (due to its aromatic structure) (Dou et al., 2016). I) BODIPY TR Methyl Ester is small lipophilic boron-dipyrromethene dye, known to occupy the interstitial space (Cooper et al., 2005; Reddy et al., 2008), possibly making it even more resistant to detergents. Alkyl-chains are blue and fixable dyes (C-F) are highlighted in the grey box.

Lipophilic dyes are prone to aggregation in physiological solutions, which leads to heterogeneous staining and self-quenching (Zimcik et al., 2007; Konno et al., 2017). Incorporating the dyes into the bilayer of liposomes can prevent aggregation and allow the dyes to more easily integrate into the cellular membranes (Blumenthal et al., 2002; Ravnic et al., 2005; Salehi et al., 2018). Especially neutral liposomes (NL) and anionic liposomes (AL) have been found to decrease DiIC18 aggregation and enhance distribution in capillaries (Konno et al., 2017). Hence, we included NL and AL to improve vessel staining in conjunction with lipid clearing.

We imaged the cerebral vasculature and the choroid plexus and evaluated the homogeneity of perfusion vessel staining before and after tissue clearing. This was to asses dye performance and compatibility with the long and harsh lipid removal and detergents during PACT (passive CLARITY technique). We used BrainFilm (Kim et al., 2018), a technique to shrink, flatten, and dry the cleared tissue, to reduce imaging and processing time, and a deep learning image segmentation tool to quantify vessel radius and dye intensity before and after PACT.

## Results

### Liposomes reduce dye aggregation and improve staining

The lipophilic dyes are highly hydrophobic and aggregate in the physiological ionic conditions. To reduce dye aggregation in the perfusate, we incorporated them into liposomes, which form small micelles around the dyes and these get integrated into the cellular membranes (MacDonald, 1990; Ertürk et al., 2011) during perfusion (Salehi et al., 2018; Li et al., 2008). We employed neutral (NL) and anionic liposomes (AL) while omitting cationic liposomes as they increase aggregation for DiIC18 (Konno et al., 2017). We examined aggregation after 30 minutes in suspension in PBS with or without liposomes.

Adding NL or AL for NR, FM 1-43FX, R18, and DiIC12, did not affect aggregation since the size and number of aggregates are similar compared to PBS (figure S1). In contrast to a previous report (Konno et al., 2017), we find that the AL resulted in a lower or equal number of aggregates compared to the NL solution (figure S1). Nevertheless, most of the fixable dyes, BODIPY, DiIC18-DS, SP-DiIC18, and CM-DiIC18, as well as DiIC18 the addition of liposomes had a reduction in aggregation.

### Dye size impacts staining specificity

To assess the effect of liposome type on the homogeneity of vessel staining and the retainment of dyes after lipid removal in PACT, we performed a series of tests. All nine dyes were coupled with either NL or AL and added to the perfusate before fixing. Brains were sliced and imaged before and after PACT with the same microscope and settings. The smallest dyes lacking alkyl-tails (i.e., Nile Red, FM 1-43FX, and BODIPY), diffused beyond the vessel wall and seeped out into the surrounding tissue, and were less suitable for perfusion-based staining. Before PACT, the small dyes have a high background with no visible signal contained in the vasculature (figure S2). The alkyl-tails are necessary for rapid membrane staining in solutions, as also observed for DiIC1(5), a DiI variant lacking the alkyl-tails (Nicola et al., 2009). For the heavier dyes with one or more alkyl-chains (i.e., R18 and DiI-dyes), homogeneity of vessel stain was related to the amount of aggregation (figure S1 with less aggregation resulting in vessel-specific signal and lower background fluorescence (figure 2A) after PACT.

**Figure 2:**
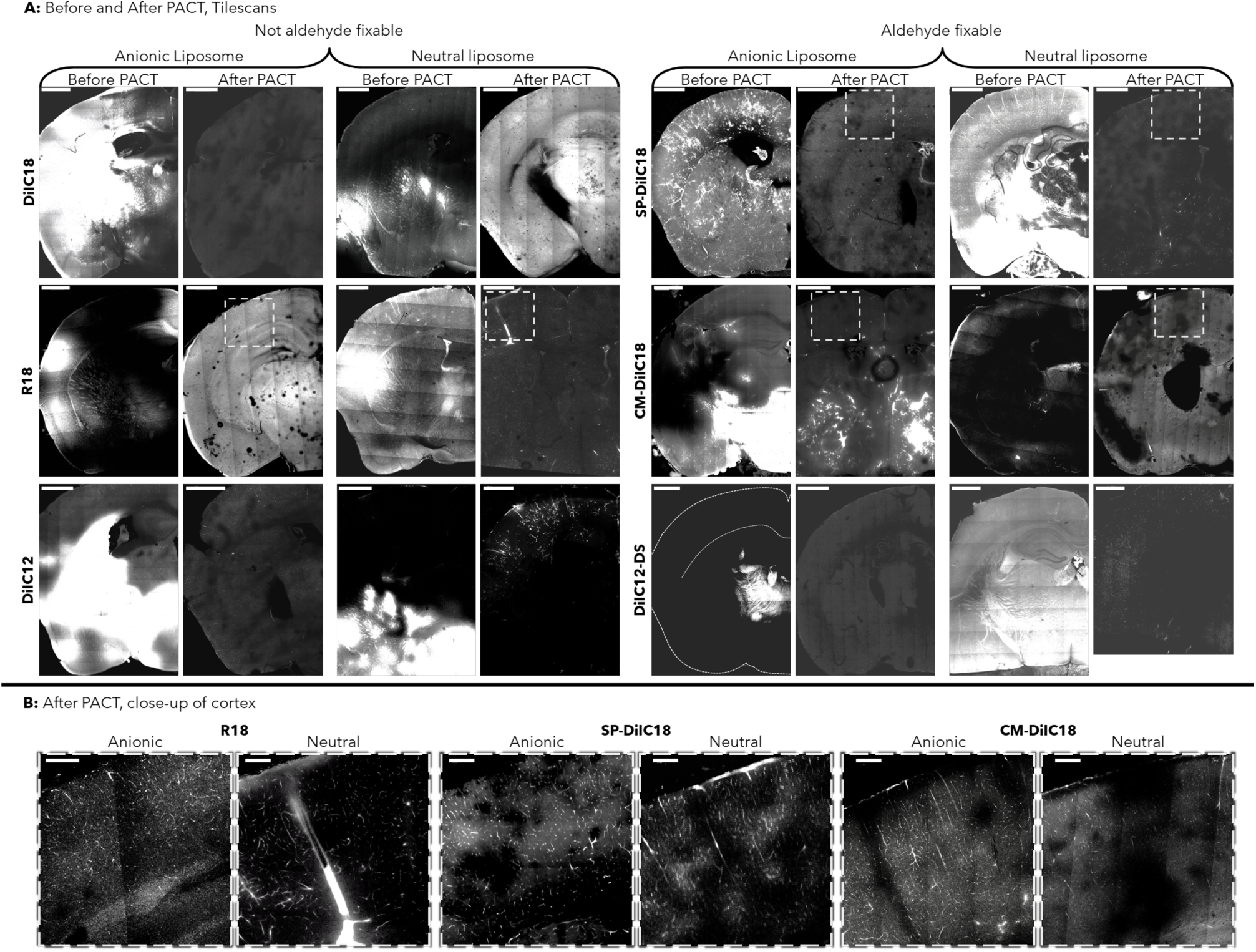
Washout of fixable and non-fixable dyes with alkyl-chains after PACT. The dyes were in anionic or neutral liposomes and delivered by perfusion before and after PACT. A) All samples had some fluorescent signal (white) after PACT. The non-fixable dyes (left) were better retained in capillaries using neutral liposomes, whereas anionic liposomes better retain the fixable dyes (right). B) Close-up of cortex with individually adjusted contrast shows clear capillaries for R18 and SP-DiIC18. Section thickness; 100 um (before PACT) and 1 mm (after PACT) coronal sections of mouse brains at bregma −1.3mm. Images were acquired and processed in a unified way. Scale bar 1 mm (A), 200 µm (B).

### Fixable dyes, CM-DiIC18, SP-DiIC18, and R18, survive PACT

Non-fixable dyes, such as DiIC18, DiIC12, and R18, in AL, were less resilient to PACT compared to utilizing NL which resulted in high vessel-specific fluorescence and low background noise before and after PACT (figure 2A).

The fixable versions of DiIC18, i.e. SP-DiIC18, and CM-DiI, combined with either liposome was present inside the vasculature after PACT. Applying anionic liposomes improved vasculature specific signal after PACT, compared to using neutral liposomes (figure 2B). The fixable analog of DiIC12, DiIC12-DS, kept most vasculature specific signal after PACT when combined with NL, but not AL (figure 2A). The dye/liposome combination of SP-DiIC/AL excelled with a homogenous stain of vessels and capillaries with nearly no background noise, both before and after PACT (figures 2A and B).

These results indicate that the alkyl-length is not crucial for obtaining a homogenous vessel stain, while fixability combined with long side chains increases PACT survival.

### Anticoagulant and vasodilator improved vessel-staining

Previous reports reports show vessel-staining is not only affect by dye aggregration, but also by dye dispersion, which is inhibited by vascular occlusion and constriction during perfusion (Ravnic et al., 2005; Weyers et al., 2012; Ghanavati et al., 2014; Salehi et al., 2018). Vasoconstriction was initially preempted by keeping perfusion fluids at 37°C. Since there was heterogeneous staining (figure 2 we added an injection of a vasodilator (sodium nitroprusside, SNP) and anticoagulant (heparin) two minutes before euthanasia.) Two dyes were selected for the optimized perfusion procedure, SP-DiIC18 and R18, since these were compatible with either AL or NL and also most cost-effective.

Injecting heparin and SNP before euthanasia resulted in increased vessel staining success for SP-DiIC18/NL and R18/NL (figure 3 In contrast, SP-DiIC18/AL vessel staining was less successful compared to the first trial (figure 2).

**Figure 3:**
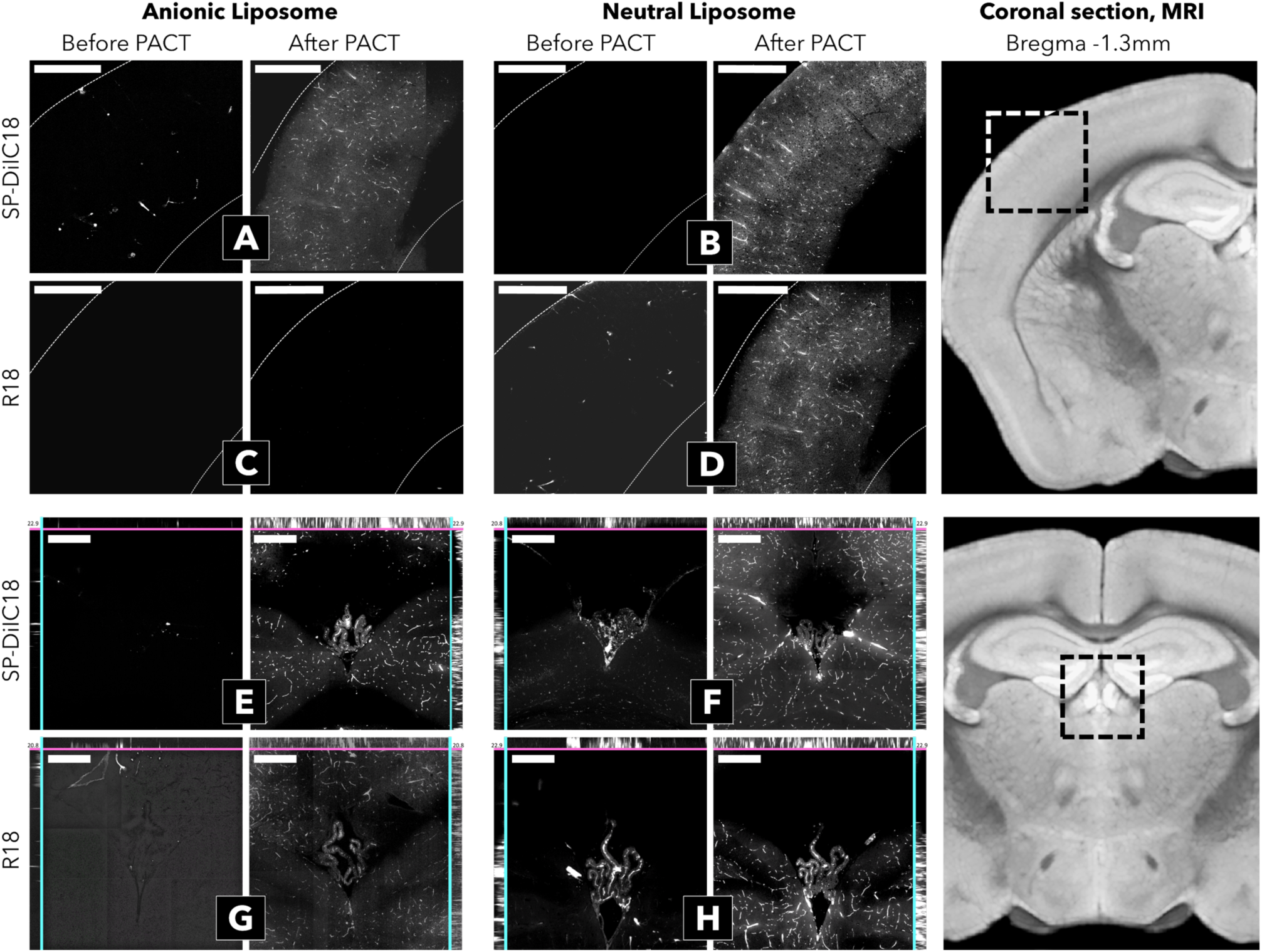
Specific liposomes, SP-DiIC18/AL and R18/NL, with vasodilators and anticlotting agents improve staining. Heparin and SNP injected before euthanasia to prevent clotting and constriction resulted in increased vessel staining success. The fluorescent in the cortical vasculature was present when using SP-DiIC18 in anionic liposomes before and after PACT (A). Whereas when using neutral liposomes, the staining was more apparent after PACT (B). R18/AL had little or no staining (C), whereas NL provided staining of the small vessels after PACT (D). Qualitatively similar results were obtained from Z-stacks of the choroid plexus. SP-DiIC18/AL (E), SP-DiIC18/NL (F), and R18/NL (H) were retained inside the lumen in smaller capillaries and larger vessels, whereas R18/AL had little staining before PACT and retained no staining after PACT (G). Images were processed in a unified way (see methods). A-D) scale 500 µm. E-H) scale 250 µm, depth 20µm. Right: MRI images indicating the selected regions (box) used with permission (Ullmann et al., 2015).

### Optimal dyes and liposomes for vessel staining

SP-DiIC18/AL resulted in homogenous vessel staining in the cortex (figure 3A)and other areas, but a low signal in the choroid plexus before PACT (figure 3E). Background noise increased after PACT, vessels, and capillaries were visible. R18/AL was predominately staining subcortical structures in the center of the brain and lacking in the cortex the choroid plexus before PACT. After PACT, the tissue showed a similar pattern of increased signal in the choroid plexus as well as an increase in background noise (figure 3F and H).

SP-DiIC18/NL and R18/NL had similar staining patterns. There was some signal from vessels in the lower or upper part of the brain and clearly defined vessels in the choroid plexus before PACT. In contrast, after PACT, the fluorescent staining was mainly located in the capillaries of the choroid plexus and an increase in visible cortical vessels (figure 3, right).

These observations indicate that combining the dyes with AL result in less staining of larger vessels, although the signal is preserved in all structures. In contrast, NL resulted in more homogenous staining of larger vessels while preserving dye retention in capillaries after PACT. Both dye combinations had an even fluorescence along Z-axis in tissue (figure 5C).

**Figure 4:**
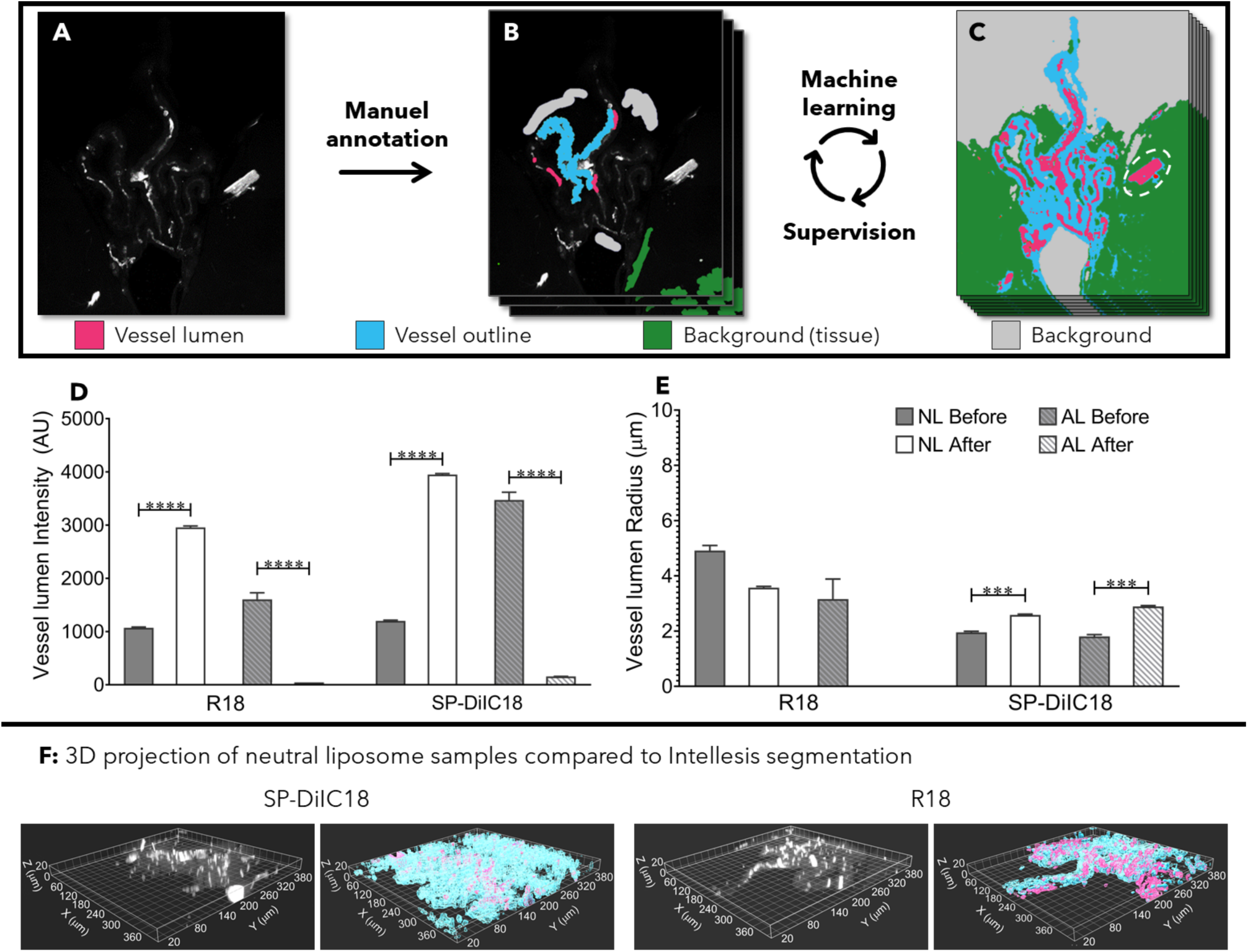
Vessel staining intensity and radius assessed using image segmentation. The median intensity in identified regions suggests SP-DiIC18 and R18 in neutral liposomes are suitable dyes in PACT samples. The deep learning image segmentation software, ZEN Intellesis (Zeiss), was used to segment the image stacks and extract vessel lumen intensity and radius. A) Sample image of the choroid plexus (dorsal 3rd ventricle) using the dye R18 and neutral liposomes. B) Supervised identification of vessel lumen (magenta), vessel outline (cyan), surrounding tissue (green), and background (grey) for training the image segmentation algorithm in Intellesis. C) Fully trained algorithm-segmentation of the entire image, with a white circle around an artefact, which was removed manually. D) Median vessel lumen intensity ± SEM from integrated z-stack of the segmented regions before and after PACT. E) Median vessel radius ± SEM (same regions as in D) R18/NL vessel radius was similar before and after PACT with a small insignificant decrease in after PACT tissue. Data was analyzed using one-way ANOVA (Kruskal-Wallis), with a multiple comparison and False Discovery rate test (Benjamin, Krieger, and Yekutieli). ****P-value<0.0001 and ***P-value<0.001. F) 3D-projections of fluorescent and surface plots of segmented vessel lumen (magenta) and outline (cyan). Z = 20 µm.

**Figure 5:**
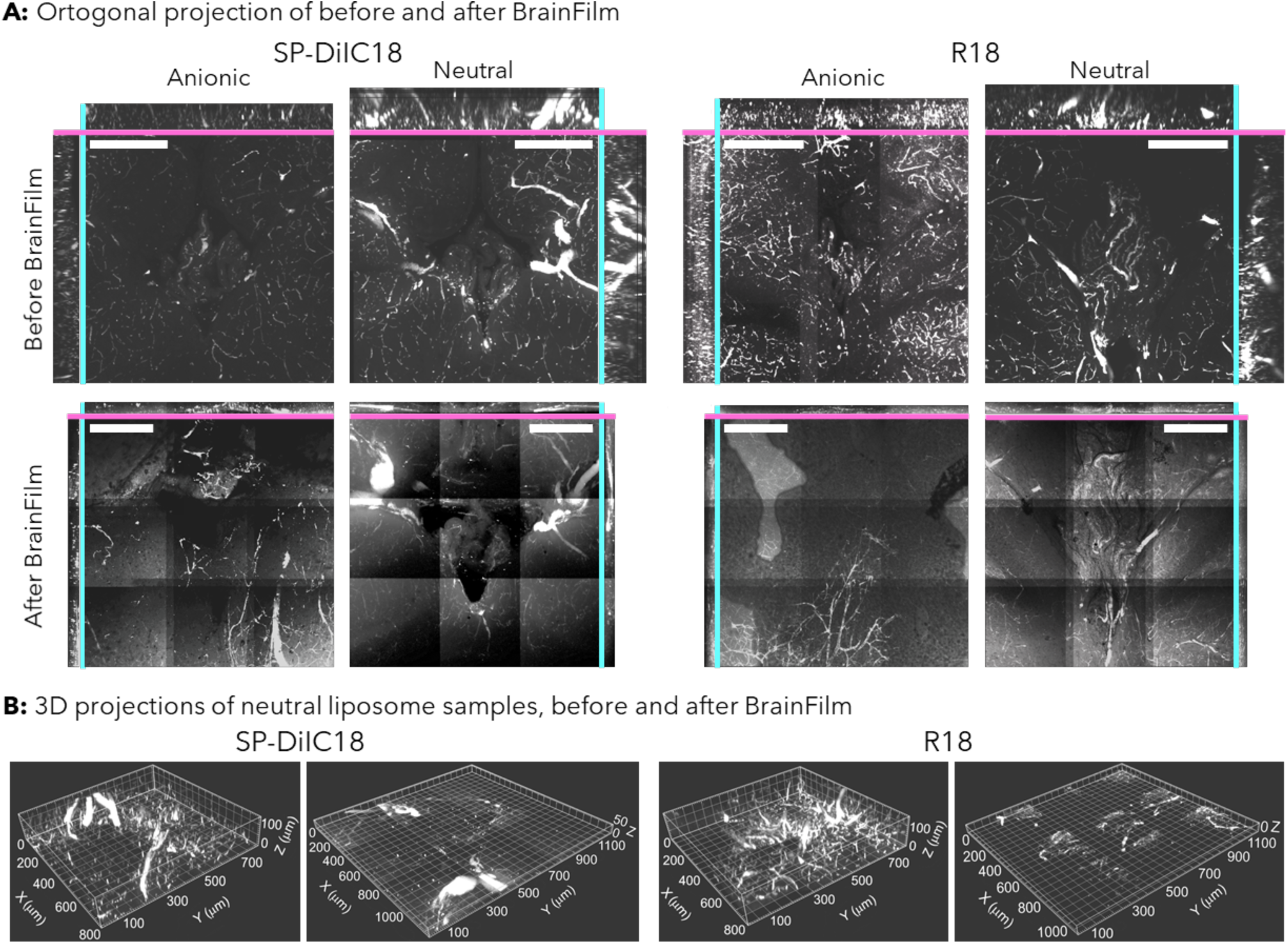
Fluorescent dyes and BrainFilm: decrease of imaging time and more background. A) The ortogonal projections of the largest possible Z-stacks achieved for each dye/liposome combination, before (top) and after (bottom) the BrainFilm process. The corresponding z-stack depth at the orthogonal cross-section shown on sides and top (cyan and magenta lines). Panels represent staining with SP-Dil18 (left) and R18 (right), with either AL or NL. B) 3D projections of SP-DiIC/NL (left) and R18/NL (right), before (right of each) and after BrainFilm (left of each). For each dye, neutral liposome seems most compatible, with the highest signal-to-noise ratio for SP-DiIC18/NL. Images were processed similarly across dye/liposome conditions. A) scale 250 µm.

### Utilizing deep learning for assessment show NL induced increase in intensity after PACT

Visual inspection of vessel staining to evaluate perfusion staining and PACT survival is widely used (Kolesová et al., 2016; Konno et al., 2017; Carrillo et al., 2018). Combining this with quantifiable measurements would be useful, and obtaining automated measurements is beneficial. Therefore, we employed a commercially available deep learning image segmentation tool (ZEN Intellesis, Zeiss). The algorithm was trained on choroid plexus images from several z-positions (figure 4A). Four different tissue labels were defined; vessel lumen, vessel outline, tissue background, and background, and these categories were manually annotated (figure 4B). After training, the automated segmentation was evaluated (figure 4C), and the fully trained model was used on the entire z-stacks of each dye. The segmentation tool was straightforward to train. Minor artifacts, were identified and removed from the dataset (figure 4C).

The intensity of R18 and SP-DiIC18 in NL was increased after PACT, whereas in AL, there was a decreased intensity (figure 4D). The radius of identified vessels was increased by 32% and 60% for SP-DiIC18 after PACT (figure 4E). Furthermore, image segmentation enabled 3D surface plots of the Z-stacks to investigate morphology better (figure 4F). In conclusion, NL increased PACT survival and prevented significant washout of R18 from smaller vessels, while image segmentation proved to be a useful tool in histology.

### SP-DiIC18 samples are compatible with BrainFilm

BrainFilm is a method that physically compresses cleared tissue, reducing sample thickness and hence the imaging time. We modified the BrainFilm procedure and reduced sample thickness from 1 mm to <0.2 mm, with an image depth of 150 µm reduced to 35 µm. A complete Z-stack of the choroid plexus was reduced to 15 minutes, compared to 1 hour before BrainFilm (figure 5A). SP-DiIC18 had the most intense and best signal to background ration with both liposomes after BrainFilm (figure 5A). Choroid plexus structure was better maintained when combined with NL for both dyes (figure 5A and B). These results show that while BrainFilm is a useful tool for decreasing imaging time, some dyes are less suitable and tissue distortion as well as an increase in background noise, is to be expected.

### Fluorescence of dyes: general observations

The short-chain DiIC12 is less hydrophobic than DiIC18. It has in vessel-painting without tissue-clearing shown better staining, likely due to being soluble slightly longer (Konno et al., 2017) and faster diffusion speed in the membrane. However, we observed that DiIC12-DS, DiIC12, and the long-chained DiIC18 showed poor retention after PACT (figure 2A), which indicates that alkyl-length is not important for dye retention after tissue clearing.

Among the fixable dyes with alkyl-tails (i.e., SP-DiIC18, CM-DiIC18, DiIC12-DS), all except DiIC12-DS performed well after PACT (figure 2A). We attribute the poor performance of DiIC12-DS not to the short alkyl-tail, but to the two directly sulfonated (DS) groups which are possibly not as aldehyde-fixable, as designed (figure 1). R18, which only has one C18 tail, performed as well as the fixable double-tailed SP-DiIC18 and CM-DiIC18 (figure 2 and 3) This could indicate the need for at least one alkyl-chain and suggest that the tertiary amine of R18 is possibly fixable.

Based on our results, we recommend using SP-DiIC18 and R18, the most cost-effective fluorescent dyes, for vessel-painting when utilizing hydrogel-based tissue clearing such as CLARITY (table 1).

**Table 1.** Performance of dyes in combination with liposomes. For the aggregation test, DiIC18, SP-DiIC18, DiIC12-DS, and BODIPY had decreased aggregation compared to PBS (+). In contrast, the rest of the dyes had no significant difference in aggregation size or number (-), CM-DiIC18 was not imaged in pure PBS. PBS. DiIC18/NL, DiIC12, CM-DiIC18, SP-DiIC18, DiIC12-DS/NL, and R18/NL all achieved vessel-specific staining (+), with the same colors having visible vessel-specific signal after PACT (PACT survival, +). DiIC18/AL, DiIC12-DS/AL, and R18/AL achieved less vessel specific and more heterogeneous staining (-). FM, NR, and BODIPY had little to no vessel-specific staining (-). BrainFilm decreased vessel-specific signal in R18 stained samples, with some specific signal remaining in R18/NL (+) compared to R18/AL (-), while the signal was sustained in SP-DiIC18 stained samples (+).

| Dye | FM 1-43FX |  | Nile Red |  | BODIPY |  | DiIC12 |  | DiIC12-DS |  | DiIC18 |  | CM-DiIC18 |  | R18 |  | SP-DiIC18 |  |
| --- | --- | --- | --- | --- | --- | --- | --- | --- | --- | --- | --- | --- | --- | --- | --- | --- | --- | --- |
| Liposome | AL | NL | AL | NL | AL | NL | AL | NL | AL | NL | AL | NL | AL | NL | AL | NL | AL | NL |
| Aggregation | - | - | - | - | + | + | - | - | + | + | + | + |  |  | - | - | + | + |
| Vessel-stain | - | - | - | - | - | - | + | + | - | + | - | + | + | + | - | + | + | + |
| PACT-survival |  |  |  |  |  |  | + | + | - | + | - | + | + | + | - |  | + | + |
| Signal depth |  |  |  |  |  |  |  |  |  |  |  |  |  |  | - |  | - | + |
| BrainFilm |  |  |  |  |  |  |  |  |  |  |  |  |  |  | - |  | + | + |
| Price/stain (USD) | 240 |  | 2.5 |  | 70 |  | 2.5 |  | 9 |  | 2.5 |  | 250 |  | 25 |  | 70 |  |

## Discussion

The vasculature of the brain is vital for the survival of neurons and investigating the structure in 3D can provide crucial insight into the origin of different pathological states. The 3D structure can be achieved post-mortem by tissue clearing, where the vessels have been perfusion-stained with fluorescent probes (Ertürk et al., 2012; Qi et al., 2019; Hsu et al., 2017; Cai et al., 2019) and serial z-stack microscopy using light sheet or confocal microscopes (Carrillo et al., 2018; Liebmann et al., 2016; Feng et al., 2017). The tissue clearing removes primarily lipids but also other molecules, including the dye for staining the vasculature, which depends on the type of dye and how well it integrates into the fixed protein fabric. The objective of the current study was to identify a lipophilic dye that does not get removed in the clearing process. We performed tests of a series of fluorescent lipophilic dyes for their retention in the vessel wall after tissue clearing. We find that SP-DiIC18, R18 and CM-DiIC18 work well even after prolonged lipid clearing, but other dyes also work.

Hydrogel-based tissue clearing techniques, such as CLARITY, can be used either with endogenously expressed fluorescent proteins or with the aid of fluorescent dyes (Jensen and Berg, 2017). Lipophilic fluorescent dyes, which are also aldehyde-fixable, are likely to provide a better fluorescent signal since these dyes are better retained in the tissue by fixing to the adjacent lipoproteins (Jensen and Berg, 2016). Here, we tested the CLARITY-compatibility of nine dyes, where some are also fixable, including DiIC variants, and found high-quality vasculature-staining (Konno et al., 2017; Salehi et al., 2018; Li et al., 2008; Ravnic et al., 2005) with most of the dyes. The dyes were added in the perfusate to stain the endothelium of vessels. To minimize the dye aggregation, which occurs in most PBS solutions, we used anionic and neutral liposomes (Konno et al., 2017). The tissue was then cleared using CLARITY, and the extent of staining was assessed with the aid of deep learning image segmentation.

### Automated image segmentation

Quantifying vessel staining using automated segmentation reduces the workload, especially compared to using tools like Fiji (Pan et al., 2016) or measuring fluorescent residue in clearing solution (Magliaro et al., 2016; Ke et al., 2013). Using the deep learning segmentation tool (Intellesis from Zeiss) we achieved reliable data collection with a further reduction in workload. An alternative to the software (intellesis) is the software AngioTool, which has been used previously in a similar manner (Salehi et al., 2018). Intellesis was developed with the intention of analyzing cell cultures and other more uniform image stacks, and here we demonstrate that it can also be applied for tissue analysis.

### Optimal dye and liposomes for vessel staining with PACT

SP-DiIC18 and R18, were compatible with both liposome solutions as the aggregate size was decreased compared to PBS. AL outperformed the NL solution with regards to aggregate size, but not when comparing vessel staining results. Visual inspection of SP-DiIC and R18, under optimal conditions, show that the staining pattern in between the dyes is very similar both before and after PACT. Using NL resulted in homogenous staining before and after PACT. In contrast, AL resulted in less staining of larger vessels before PACT with some signal remaining in all structures and increased background noise after PACT, compared to NL. This could be because AL is less compatible with the surface charge of endothelial cells (Konno et al., 2017).

Data from segmented images showed an increase in intensity for SP-DiIC18/NL and R18/NL after PACT, which could be due to the increased transparency after PACT, allowing an increased amount of fluorescence to escape from the lower layers. This would also result in more well-defined smaller vessels, which was observed by visual inspection of the tissue after PACT. Vessel radius remained consistent in R18/NL tissue sections before and after PACT, whereas in the SP-DiIC18/AL and SP-DiIC18/NL staining, there was a significant increase of vessel radius after PACT. This indicates that R18/NL is more able to retain in the smaller and larger vessels, whereas SP-DiIC18 is washed out of smaller vessels or bleed out into the vessel wall, making vessel size less consistent.

### Compatibility with other tissue clearing and processing techniques

We used BrainFilm to flatten and dry the tissue for faster imaging and processing. We replaced cellophane with glass slides, because we believe the cellophane distorted our images due to refractive index mismatch with our protocol. SP-DiIC18 was better than R18 when using BrainFilm since R18 gave more background fluorescence with less stained vessels (figure 5). The decrease in signal could be self- or concentration-quenching, which R18 is known for, due to increased dye concentration due to compression and drying. The decrease in signal to background ratio could be due to the volume compression and hence increase in fluorescent noise per volume.

We used PACT, which relies on prolonged incubation, agitation, and high detergent concentrations at 37°C, and BrainFilm is harsh with drying and further impacts the fluorescence signal setting high demands of dyes. CM-DiIC18, SP-DiC18 and R18 performed well in PACT. SP-DiC18 outperformed R18 in BrainFilm. CM-DiIC18 e.g. unlike DiI is retained in cells throughout fixation, permeabilization, and paraffin embedding procedures. Acetone treatment of SP-DiC18 stained cells has according to the manfucaturer even enhanced fluorescence, which makes SP-DiC18 and other sulfonated DiI-dyes ideal for tissue clearing which requires dehydration, clearing, and lipid extraction.

Hence, we predict that SP-DiIC18 and CM-DiIC18, tested under several harsh conditions here is also compatible with other clearing methods based on steps of dehydration, lipid removal, organic solvents, detergents or electrophoresis as 3DISCO, CUBIC, Expansion Microscopy, TDE, PEGASOS and ethyl-cinnamate. Moreover, perfusion based-vessel painting dyes may also be useful in examining the vasculature and lymphatic system of other organs and animals, or the xylem and phloem of plant tissues; that require CLARITY clearing with electrophoresis or the addition of enzyme digestions.

## Conclusion

Of the nine tested dyes, SP-DiIC18, and R18 are the most cost-effective and suitable dyes for vessel painting in the context of hydrogel-based tissue clearing, such as CLARITY. Both can be used together with liposomes to increase efficiency further. Anionic and neutral liposome (NL) solutions were both applicable, although the NL solution resulted in less variation across vessel painting trials. When using BrainFilm, R18 lost considerable fluorescence, making SP-DiIC18 the most suitable dye for this and other methods requring dehydration. Automated image segmentation (Intellisis from Zeiss) can be applied and help speed up the processing and quantification.

## Limitations of study

The current study aims to test various fluorescent lipophilic dyes for their compatibility with hydrogel-based tissue clearing, such as CLARITY, in the context of staining the vasculature. There are many parameters to alter and different strategies to pursue, and therefore it is difficult to find the optimal conditions for vessel staining. For instance, The compatibility of perfusion with fluorescent or biotinylated lectins (Robertson et al., 2015) with CLARITY was not included in our study. Similarly, BODIPY conjugated to lipid and cholesterol analogs, and would likely behave similar to the alkyl-tailed DiI-dyes, but were omitted due to their high price. Furthermore, it would have been more appropriate to use a light sheet microscope, which is designed for 3D imaging and reconstruction, rather than a confocal microscope. Nevertheless, we did not have access to a light sheet microscope, which is the ideal method for large volume imaging of cleared tissue (Jensen and Berg, 2017).

## Methods

Our experiments involve *in vitro* and *in vivo* experiments on 28 male mice (8- to 14-week-old C57BL/6J,). All experiments were approved by the Department of Experimental Medicine and performed according to procedures laid out by the Danish Ministry of Justice and the Danish National Committee for Ethics in Animal Research and guidelines of the Council of the European Union (86/609/EEC).

### Liposomes and Dyes

The anionic liposome (AL) solution was freshly made of 5.5mg/mL L-α-Lecithin (Merck Millipore, 524617) and cholesterol 1.18mg/mL (Merck Millipore, 228111) in ethanol, and the neutral liposomes (NL) was resolubilized according to the manufacturer (EL-01-N, NOF Corporation) in phosphate-buffered saline (PBS), before the experimental procedure utilizing sonication. Larger liposomes were removed using a filter (0.2 µm filter) before dyes were added to the solutions followed by 1 minute of sonication, which will place dyes into the liposomes. The final concentration of the dyes, Nile Red (NR), FM 1-43FX, BODIPY, DiIC18, CM-DiIC18, SP-DiIC18, Octadecyl Rhodamine B (R18), DiIC12 (Thermo Fisher Scientific) and DiIC12-DS (ATT Bioquest), was kept constant at 50 µM throughout the experimentation and the lipid concentration at approximately 1mg/mL, as in (Konno et al., 2017). To asses aggregation, each dye was suspended in a liposome solution or PBS with 4% glucose, and two 20 µL drops of the solutions were placed on a glass slide and imaged after 30 minutes.

### Dye Perfusion

For *in vivo* vessel-painting, we deeply anesthetized mice with intraperitoneally injected pentobarbital, surgically opened the chest and transcardially perfused 10 ml of dye/liposome solution (30-37°C) followed by 10 ml of fixative (4% PFA, 5-10 °C). The vessel painting process left the feet and nose slightly purple or blue if successful. Following PFA-perfusion, organs were removed and stored in fixative overnight.

In the last experiment with only SP-DiIC18 and R18 an intraperitoneal injection of 0.15 mL sodium nitroprusside (125 µg/mL; 0.75 mg/kg) and heparin (333 units/mL; 2000 units/kg) was added two minutes before euthanasia.

### PACT

We used the Passive CLARITY technique (PACT) (Yang et al., 2014) and increased the pH of the 8% sodium dodecyl sulfate (SDS) solution to 8.5 (Treweek et al., 2015). Briefly, overnight PFA-fixed tissue was rinsed in PBS and incubated overnight at 5oC with 3mL degassed hydrogel monomer solution. Tissue was polymerized in solution at 37°C for 3-4 hours, and tissue was extracted from the gel with Kimwipes (Kimwipes, Sigma Aldrich Z188956). Tissue was embedded in 4% low-melting agarose in PBS (SeaPlaque, Lonza), mounted on a vibratome with cold PBS. 1 mm coronal sections between −0.50 and −3.00 mm Bregma were used. Tissue-clearing was in 8% SDS at 37°C until fully cleared (6 days), with SDS changed every second day to speed up the process. After PACT, tissue was washed five times (10 minutes) in PBS at room temperature, and dried gently with Kimwipes and transferred to a refractive index matching (RIM) solution of 63% (v/v) 2,2′-thiodiethanol (TDE, VWR international) in 10 mM PBS, pH adjusted to 7.5 with NaOH. The tissue was incubated overnight in the RIM at room temperature, RIM was changed, and the tissue was stored until imaging (maximum one week after RIM change).

### Modified BrainFilm

From each dye and liposome condition, a 1 mm coronal brain slice that was cleared with PACT and incubated in RIM was treated with a modified version of the BrainFilm (Kim et al., 2018) to reduce the thickness to 1/10 of the original for less labor-intensive and faster microscopy. Briefly, slices were washed in PBS (5 minutes), and the kit was assembled using cover glass slides (Menzel-Gläser), and filter paper (0.20 µm pore size) inside an acrylic mold (gel drying kit, Sigma Aldrich). The BrainFilm process comprised of increasing pressure to a total of 1.5 kg on top of the mold every 2-3 hours while the mold was at room temperature, increasing ambient temperature to 45oC reduced the processing time to half. The mold clips were then applied, and the amount of filter paper was increased every 40 minutes (3-4 times), before the mold was left overnight, or to completely dry. Finally, the two pieces of coverglass were sealed with tape.

### Microscopy

For the assessment of aggregation and estimation of PACT survival and optimization, imaging was done on a widefield microscope (Zeiss AxioImager Z1, AxioCam 506 mono, or MRc5 camera) with EC Plan-Neofluar 10x/NA0.3 and 20x/NA0.5.

While imaging cleared samples, the tissue was placed in a chamber designed for live-cell imaging (POC R2, PeCon) consisting of two 0.17 mm glass slides with silicon spacers to prevent compression. The tissue was sandwiched between the glass slides while in RIM and gently held in place without compression by the silicon spacers. All tile and z-stack images of *in vivo* vessel-painting were done using a confocal LSM 710 (Zeiss) with either the EC Plan-Neofluar, 10x/NA 0.3 objective (5.20 mm working distance) or the Plan-Apochromat, 20x/0.8 objective (0.55 mm working distance). Excitation sources were a 25 mW Argon laser at 488 nm and 514 nm, and a 20 mW Solid State at 561 nm. For the first experiments, the laser power was kept constant at approximately 4% power in between dyes, before and after PACT, for the last experiment the specific laser power was adjusted for each dye before PACT, then measured using an optical power and energy meter (PM200, Thorlabs), and the same laser power was used after PACT. Images were acquired with two times frame–averaging, 0.98-1.2 Airy unit, 10% overlap in XY-planes.

The stitching of overlapping images was performed in Zen 3.1 (Zeiss) or Huygens Professional version 19.10 (Scientific Volume Imaging). The maximum intensity and 3D projections were made in Imaris Viewer 9.5.1, which were then optimized in Paint.net 4.2.10 with the same parameters for the individually compared pictures.

### Image Visualization and Deep Learning Segmentation

Quantifiable measures were obtained using a deep learning image segmentation software, (Zen Intellesis, Zeiss) on 16-bit images. Intellesis utilizes a supervised deep learning algorithm to identify Regions of Interest (ROI). During this first step, four different ROI labels were manually defined; vessel lumen, vessel outline, tissue background, and background (figure 4A). The model was trained on three images at different Z-positions for two different dyes, one with high intensity and one with low intensity. Following training, the model segments an entire image, and training is repeated if the ROI detection and labeling are unsatisfactory compared to manual detection. Since the fully trained model bases detection on intensity, structure, texture, and edge detection, two differently trained models are used in the ROI quantification, one based on images before PACT and one based on after PACT images. The trained model was used to measure vessel lumen intensity, radius, area as well as background intensity for the entire dataset. Artifacts and falsely identified vessels were manually identified and removed, using Intellesis-analyzed pictures created by the model.

### Statistical Analysis

Data was analyzed using statistical software (Prism 8, Graphpad), and results are presented as mean ± SEM, with all data points. Two different comparisons were made, and both statistical tests were used in analyzing the results. For each dye/lipid solution data before and after PACT was compared using a two-tailed, unpaired t-test (Mann-Whitney) was performed. A one-way ANOVA (Kruskal-Wallis), with a multiple comparison and False Discovery rate test (Benjamin, Krieger, and Yekutieli), was performed on data sets containing all data points for all dye/lipid solutions, before and after PACT. No assumption of Gaussian distribution was used in all analyses, and a P-value <0.05 was considered statistically significant.

## Acknowledgements

We thank the Core Facility for Integrated Microscopy, Faculty of Health and Medical Sciences, University of Copenhagen, and their assistance provided by Thomas Braunstein and Pablo Hernandez-Varas. This research was supported by the Independent research fund Denmark, Carlsberg Foundation (R. W. B.), Lundbeck Foundation (R.W.B.), and A.P. Møller og Hustru Chastine Mc-Kinney Møllers Fond til almene Formaal (R.W.B.).

## Author contributions statement

B.L.E.H., R.W.B, and K.H.R.J conceived the experiments, B.L.E.H. conducted the experiments. All authors analysed the results. B.L.E.H. and R.W.B. wrote the initial manuscript.

## Supplemental Figures

**Figure S1:**
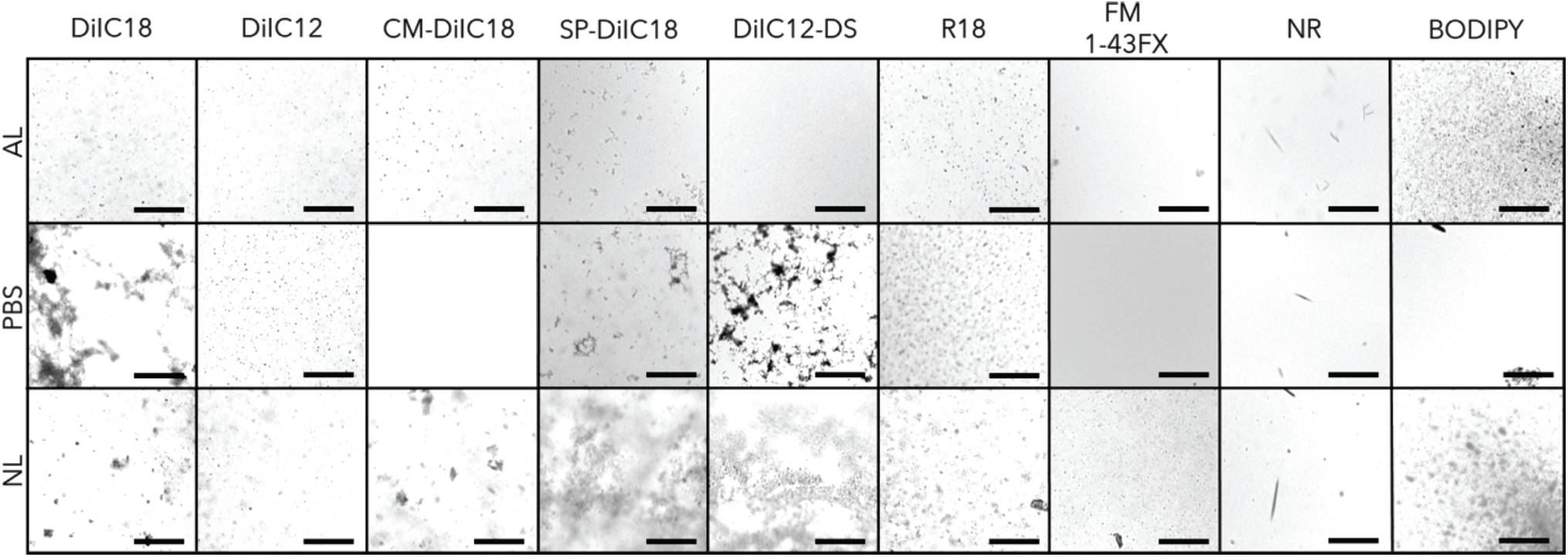
Dye aggregation after 30 minutes in PBS-glucose or with added neutral (NL) or anionic liposomes (AL). Adding liposomes reduced aggregation for most dyes, especially BODIPY, DiIC18-DS, SP-DiIC18, and CM-DiIC18, benefited from the addition of AL with a decrease in aggregate size compared to NL and PBS. NR, FM 1-43FX, R18, and DiIC12 aggregation size and number are similar across the three conditions. Scalebar 200 µm.

**Figure S2:**
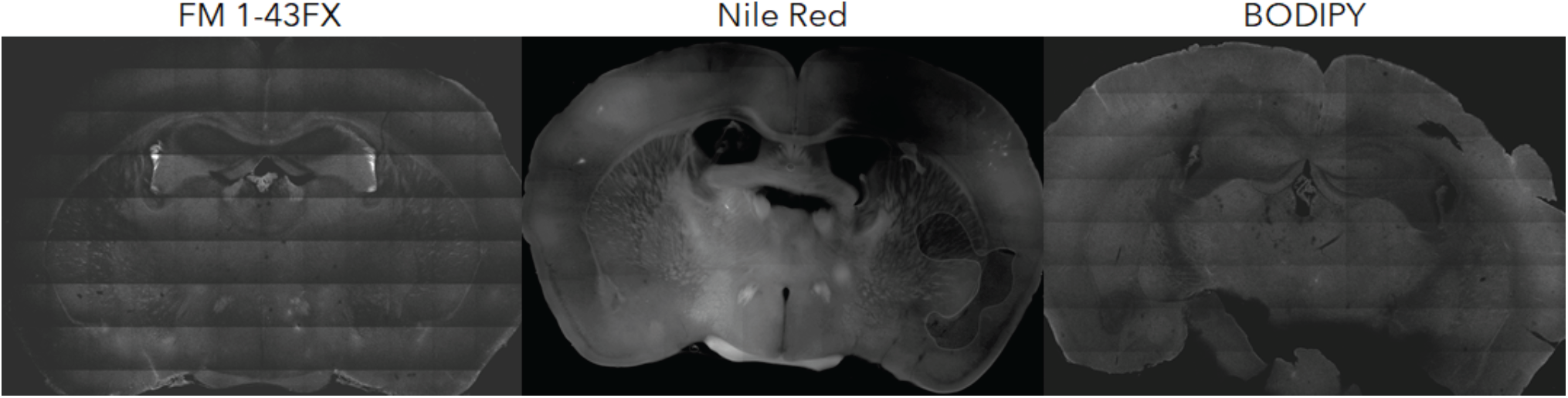
Small dyes performed poorly. Before PACT samples containing FM 1-43FX (left), Nile Red (middle), and BODIPY (right) were evaluated. The dyes penetrated into the surrounding tissue giving a large background signal with no apparent specificity for endothelial cell walls. For this reason, these dyes were not used further in the study.

